# The role of CLV signalling in the negative regulation of mycorrhizal colonisation and nitrogen response of tomato

**DOI:** 10.1101/2020.07.03.185991

**Authors:** Chenglei Wang, Karen Velandia, Choon-Tak Kwon, Kate E. Wulf, David S. Nichols, James B. Reid, Eloise Foo

## Abstract

Plants form mutualistic nutrient acquiring symbioses with microbes, including arbuscular mycorrhizal fungi. The formation of these symbioses is costly and plants employ a negative feedback loop termed autoregulation of mycorrhizae (AOM) to limit arbuscular mycorrhizae (AM) formation. We provide evidence for the role of one leucine-rich-repeat receptor like kinase (FAB), a hydroxyproline *O*-arabinosyltransferase enzyme (FIN) and additional evidence for one receptor like protein (*Sl*CLV2) in the negative regulation of AM formation in tomato. Reciprocal grafting experiments suggest that the *FAB* gene acts locally in the root, while the *SlCLV2* gene may act in both the root and the shoot. External nutrients including phosphate and nitrate can also strongly suppress AM formation. We found that FAB and FIN are required for nitrate suppression of AM but are not required for the powerful suppression of AM colonisation by phosphate. This parallels some of the roles of legume homologs in the autoregulation of the more recently evolved symbioses with nitrogen-fixing bacteria leading to nodulation. This deep homology in the symbiotic role of these genes suggests that in addition to the early signalling events that lead to the establishment of AM and nodulation, the autoregulation pathway might also be considered part of the common symbiotic toolkit that enabled plants to form beneficial symbioses.

**Highlight:** We describe the role of CLV signalling elements in the negative regulation of arbuscular mycorrhizal symbioses of tomato, including influencing nitrate but not phosphate suppression of mycorrhizal colonisation.

## Introduction

Plants can form beneficial symbiotic relationships with a variety of soil microbes. The symbiosis with arbuscular mycorrhizal (AM) fungi is ancient and widespread, occurring in over 80% of terrestrial plants and supplying plants with previously inaccessible nutrients and enhancing stress tolerance (Martin *et al.*, 2017; Pozo *et al.*, 2010; Smith and Read, 2010). Nodulation is the symbiosis between nitrogen fixing rhizobial bacteria and predominantly legumes and is thought to have evolved in part by recruiting part of the pre-existing AM signalling pathway, including the common symbiotic pathway that enables initial communication and symbiotic establishment (Delaux *et al.*, 2015; Radhakrishnan *et al.*, 2020). As the formation of these symbioses are energetically costly (Douds *et al.*, 2000; Schulze *et al.*, 1999), the plant must tightly control the ultimate extent of the symbioses. Autoregulation enables plants to limit the extent of symbioses via a negative feedback loop, reviewed by Wang *et al.* (2018). Studies in legumes indicate at least some of the genetic elements in autoregulation of nodulation (AON) overlap with elements of autoregulation of mycorrhizae (AOM) but have also highlighted some important differences (e.g. Catford, 2003; Foo *et al.*, 2016; Müller *et al.*, 2019). However, until now our genetic understanding of the genes and signals that limit mycorrhizal colonisation has been largely limited to legumes. In this paper, we use the model non-legume tomato to provide fundamental information of this key genetic program.

Our understanding of AON is relatively advanced from studies in legumes including *Medicago truncatula, Lotus japonicus,* soybean (*Glycine max*) and pea (*Pisum sativum*). The systemic AON feedback loop begins with root events associated with nodulation inducing a specific subset of CLE peptides, some of which are tri-arabinosylated by a hydroxyproline *O-* arabinosyltransferase enzyme (*Ps*NOD3, *Mt*RDN1 and *Lj*PLENTY) (Hastwell *et al.*, 2018; Imin *et al.*, 2018; Kassaw *et al.*, 2017; Okamoto *et al.*, 2013;

Yoro *et al.*, 2019). In *L. japonicus* CLE peptides are translocated to the shoot and it is clear that perception by shoot acting receptor complex(es) occurs across several species. Key players in this perception system are leucine-rich-repeat receptor like kinases (LRR-RLK) including CLAVATA1 (CLV1) - like (*Gm*NARK, *Lj*HAR1, *Mt*SUNN and *Ps*SYM29) (Krusell *et al.*, 2002; Nishimura *et al.*, 2002; Schnabel *et al.*, 2005; Searle *et al.*, 2003), KLV (Miyazawa *et al.*, 2010; Oka-Kira *et al.*, 2005), the pseudo-kinase CRN (Crook *et al.*, 2016) as well as the leucine-rich-repeat receptor like protein CLV2 (Krusell *et al.*, 2011). The perception of the CLE signal(s) activates a shoot-derived signal(s) that is transported to the root and inhibits further nodule formation (Lin *et al.*, 2010; Okamoto *et al.*, 2009; Sasaki *et al.*, 2014). Key downstream root acting players include the kelch repeat-containing F-box protein TML (Gautrat *et al.*, 2019; Magori *et al.*, 2009; Takahara *et al.*, 2013), and the transcriptional regulation of nod factor receptors (Gautrat *et al.*, 2019). A shoot-derived systemic miRNA, miR2111, maintains susceptible status in non-nodulated roots and can suppress subsequent nodulation by activating *TML* in a *HAR1* dependent manner (Tsikou *et al.*, 2018). Indeed, miR2111 appears to be a central player in both negative and positive regulation of nodulation as it also acts downstream of the carboxyl-terminally encoded peptide (CEP) perception system via the CRA2 receptor that positively regulates nodule formation (Gautrat *et al.*, 2020; Imin *et al.*, 2013; Laffont *et al.*, 2019). It is also important to note that split-root studies with *sunn* and *rdn1* mutants in *M. truncatula* suggest that there are likely to be multiple systemic regulatory pathways controlling nodulation (Kassaw *et al.*, 2015).

Disruption in elements of the AON pathway lead to an excess nodulation (super/hypernodulation) phenotype and early studies revealed that the *clv1-like* mutants in legumes (*Mtsunn, Ljhar1, Gmnark* and *Pssym29*) also developed supermycorrhizal phenotypes (Morandi *et al.*, 2000; Sakamoto and Nohara, 2009; Solaiman *et al.*, 2000), highlighting the importance of this receptor for both AON and AOM. Recently, the *rdn1* mutant of *M. truncatula* has also been reported to display elevated AM colonisation (Karlo *et al.*, 2020). This overlap is consistent with elegant studies in legumes and non-legumes that revealed rhizobium and/or nodulation can suppress mycorrhizal development and vice versa (Catford, 2003; Khaosaad *et al.*, 2010; Sakamoto *et al.*, 2013). In contrast to this conservation, nodulation and mycorrhizae induce the expression of a specific sub-set of CLE peptides (de Bang *et al.*, 2017a; Handa *et al.*, 2015; Karlo *et al.*, 2020; Müller *et al.*, 2019). Indeed, recent studies in *M. truncatula* have established that specific CLE peptides suppress the formation of AM (Karlo *et al.*, 2020; Müller *et al.*, 2019). Overexpression of *MtCLE53* and *MtCLE33* led to significantly reduced mycorrhizal colonisation compared with the control construct, and this suppression was dependant on the *CLV1-like* gene *SUNN* and *RDN1.* A role for tri-arabinosylation in activation of *MtCLE53* was supported by the fact that overexpression of a modified version of *MtCLE53* that may be unable to be tri-arabinosylated did not influence AM (Karlo *et al.*, 2020). In contrast to the important role for *MtCLE53,* overexpression of the nodulation induced *CLE, MtCLE13,* did not suppress AM colonisation (Müller *et al.*, 2019). Intriguingly, AM fungi themselves can also produce CLE peptides that appear to promote colonisation (Le Marquer *et al.*, 2018). The genetic components of AOM in non-legumes is only now emerging, with roles for *CLV1-like* genes and *CLV2* suggested by the elevated AM colonisation observed in the *Brachypodium distachyon clv1-like* mutant *fon1-1* and transgenic lines of tomato disrupted in *CLV2* respectively (Müller *et al.*, 2019; Wang *et al.*, 2018).

Plants strongly regulate symbioses in response to nutrient availability. High phosphorus supply suppresses AM formation across species, while it promotes nodulation (e.g. Breuillin *et al.*, 2010; Foo, 2017). Nitrogen supply strongly suppresses nodulation (e.g. van Noorden *et al.*, 2016) and in some species has also been observed to suppress AM, although neutral and positive effects of nitrogen on AM have also been reported (Lim et al., 2014; Liu et al., 2012; Bonneau et al., 2013; Corrêa et al., 2014; Nouri et al., 2014). Indeed, mycorrhizal-induced ammonium and nitrate plant transporters have been identified (e.g. Guether *et al.*, 2009; Wang *et al.*, 2020). There is genetic evidence that nitrogen, and possibly phosphorous, interact with elements of the AON pathway to regulate nodulation. For example, plant mutants disrupted in the CLV1-like protein, KLV, and RDN1 in some species display reduced sensitivity to nitrate suppression of nodulation (Carroll *et al.*, 1985;

Jacobsen and Feenstra, 1984; Lim *et al.*, 2011; Oka-Kira *et al.*, 2005; Schnabel *et al.*, 2005; Searle *et al.*, 2003). Further, the pea *Psnark (clv1-like*) mutant does not suppress nodulation under low phosphate (Foo *et al.*, 2013a). In addition, CLE peptides whose expression responds to altered nutrients such as nitrate and phosphorous have been characterized in legume and non-legume systems and regulate a variety of nutrient responses, including legume nodulation (Araya *et al.*, 2014; de Bang *et al.*, 2017b; Karlo *et al.*, 2020; Müller *et al.*, 2019; Okamoto *et al.*, 2009). However, the role of these elements in regulating AM in response to nutrient status is underexplored. In pea, *M. truncatula* and soybean, although *CLV1-like* genes *PsSYM29, MtSUNN* and *GmNARK* are required to negatively regulate AM formation, these genes do not appear to be required to do this in response to phosphate as *Pssym29, Mtsunn* and *Gmnark* mutants still suppress AM under high phosphate (Foo *et al.*, 2013a; Müller *et al.*, 2019; Wyss *et al.*, 1990). However, the possibility that nitrogen influences AM formation via this pathway has not yet been addressed.

Elements downstream of AOM are still unclear. One potential player that has been proposed is strigolactone, the root exuded plant hormone that can promote establishment of mycorrhizal symbiosis by promoting arbuscular mycorrhizal fungal spore gemination and hyphal branching (Akiyama *et al.*, 2005; Besserer *et al.*, 2006). This is based on the observation that in *M. truncatula* overexpression of *MtCLE53* or *MtCLE33* down-regulated strigolactone biosynthesis and that the low colonisation rates of these lines could be elevated with strigolactone application (Müller *et al.*, 2019). However, strigolactone levels were not elevated in the *sunn* mutant of *M. truncatula* or the pea *sym29, clv2* or *nod3* mutants and double mutant studies in pea clearly indicate strigolactones do not act downstream of AON (Müller et al., 2019, Foo et al., 2014). Given the potent role for strigolactones in the up-steam establishment of AM symbioses it is difficult to establish if strigolactones also act downstream of AOM.

Tomato mutants disrupted in the *CLV1-like* gene (*fab), CLV2* gene (*Slclv2*) and *RDN1-like* gene (*fin*) are available and have been previously characterised for their role in shoot apical meristem identity and root development (Xu *et al.*, 2015). Previous studies also suggested a role for *SlCLV2* in the negative regulation of AM symbioses (Wang *et al.*, 2018). In this paper, the hypothesis that *FAB* and *FIN* play a role in the regulation of AM development of non-legume tomato was examined, and the hypothesis that nitrate and phosphate act through these genes to regulate AM was tested. The *FAB* and *FIN* genes are shown to exert a negative influence on AM formation and are required for the nitrate suppression of AM in tomato. However, they do not influence the strong suppression of AM formation by phosphate. In contrast to the shoot acting role of *CLV1-* and *CLV2-like* genes in legumes in regulating symbioses, we found only limited evidence that *FAB* and *SlCLV2* act outside root tissues to influence AM.

## Materials and Methods

### Plant materials

One study was conducted with the tomato *Solanum lycopersicum* wild type (WT) cv. Money Maker. All other experiments employed the tomato WT cv. M82, and mutants on this background, *fab, fin-n2326* and *fin-e4489* and the CRISPR generated mutant *Slclv2-2* (Xu *et al.*, 2015). The *fab* mutant carries a single base pair substitution resulting in an alanine to valine substitution in the kinase domain, the *fin-n2326* mutant has a large sequence deletion that results in the absence of transcripts and the *fin-e4489* mutant has a 1bp missense mutation that results in a premature stop codon (Xu *et al.*, 2015). Mutations introduced in *Slclv2-2* are outlined in Suppl Fig. S1 at *JXB* online.

### Growth conditions

Tomato seeds were germinated in potting mix and transplanted two weeks after, sowing into 2L pots containing a 1:1 mixture of vermiculite and gravel (plus inoculum), topped with vermiculite. Unless otherwise stated, plants were grown under glasshouse conditions (18 h photoperiod). For experiments using tomato *clv2* mutants, the plants were grown in controlled glasshouse (25 °C day/20 °C night, 18 h photoperiod). Unless otherwise stated, tomato plants were supplied with 75ml/pot modified Long Ashton nutrient solutions (Hewitt, 1966) containing 5 mM KNO_3_ and 0.5 mM NaH_2_PO_4_ twice a week (1 - 3 weeks after transplanting) and three times a week (from 3 weeks after transplanting).

Inoculum for mycorrhizal experiments was live corn pot culture originally inoculated with spores of *Rhizophagus irregularis* (INOQ Advantage, INOQ GMBH, Germany), grown under glasshouse conditions that received modified Long Ashton nutrient solution containing 3.7mM KNO_3_ and 0.05mM NaH_2_PO_4_ once a week. The inoculum contained colonised root segments, external hyphae and spores. For standard experiments, the growth substrate (80%) was mixed with corn pot culture (20%). For high dose of inoculum treatments, inoculum was increased to 40% corn pot culture.

For grafting experiments, wedge grafts were performed in the hypocotyl three days after transplantation of rootstocks and the grafts maintained in a humid environment for approx. 5 - 7 days and then gradually reintroduced to ambient conditions.

### Root staining and scoring

Tomato plants were harvested 6-8 weeks after transplanting. The root and shoot were separated, fresh weight recorded and the tomato roots were cut into 1-1.5 cm segments, except for hyphopodia measurements where whole roots were gently removed and placed in nylon Biopsy Bags (Thermo Fisher Scientific, USA) inside tissue processing cassettes (Thermo Fisher Scientific, USA).

Unless otherwise noted the ink and vinegar method was used for mycorrhizal staining (Vierheilig *et al.*, 1998). Mycorrhizal colonisation of roots was scored according to McGonigle *et al.* (1990), where 150 intersects were observed from 25 root segments per plant. The presence of arbuscules, vesicles and intraradical hyphae at each intersect was scored separately. The total colonization of mycorrhizae was calculated as the percentage of intersects that have presence of any fungal structures and arbuscule frequency was calculated from the percentage of intersects that contained arbuscules. For the hyphopodia experiment, cassettes containing root samples were covered with 5% KOH at 58 °C overnight, rinsed with water and 3.5% HCl, and then stained in 0.05% trypan blue lactoglycerol solution for 12 h at 58 °C. The number of hyphopodia was scored on 15 root segments per plant and is presented as the total number of hyphopodia per cm of root length.

### Nitrate and phosphorous influence on AM

For the nitrate experiments, plants were grown in the presence of mycorrhizal inoculum and supplied with modified LANS with 0.5mM NaH_2_PO_4_ and various concentrations of KNO_3_ (ranging from 0.625mM to 10mM for examining N impact on WT). Two N concentration (0.625mM and 10mM) were selected for examining N impact on mycorrhizal colonization in mutants. For the phosphate experiments, plants were grown in the presence of mycorrhizal inoculum and supplied with modified LANS with 5mM KNO_3_ and two concentrations of NaH_2_PO_4_ (0.05 and 5mM).

### Strigolactone extraction and quantification

Tomato plants for strigolactone analysis received 2.5mM KNO_3_ and 0.5mM NaH_2_PO_4_ nutrient solution. Root exudate was collected from individual plants and strigolactones extracted and measured by UPLC/MS-MS as outlined in (Foo and Davies, 2011). Strigolactone standards, [6’-^2^H_1_]-orobanchol, [6’-^2^H_1_]-orobanchyl acetate, [6’-^2^H] 5-deoxystrigol and [6’-^2^H_1_]-fabacyl acetate, were added to each sample solution as internal standards. As there is no labelled solanacol standard available, an un-labled solanacol sample (kindly provided by A/Prof Chris McErlean and Dr Bart Janssen) was run as an external control and after initial analysis, all samples were spiked with solanocal and re-run to ensure solanacol could be detected in the sample matrix. For solanacol, transitions monitored were 343 > 97, 343 > 183 and 343 > 228 and other for other strigolactones the ions monitored were as reported previously (Foo and Davies, 2011). The endogenous strigolactone levels were calculated from the ratio of endogenous to standard peak areas per gram root fresh weight.

### Phylogenetic analysis

The full length amino acid sequence of CLV1, CLV2 and RDN1 related proteins was used for phylogenetic analyses. The multiple sequence alignment was generated using the Muscle algorithm (Edgar, 2004). The phylogenetic tree was constructed using the Maximum Likelihood method based on the Whelan And Goldman + Freq. model (Whelan and Goldman, 2001). The trees with the highest log likelihood are shown. Initial tree(s) for the heuristic search were obtained automatically by applying Neighbor-Join and BioNJ algorithms to a matrix of pairwise distances estimated using a JTT model, and then selecting the topology with superior log likelihood value. A discrete Gamma distribution was used to model evolutionary rate differences among sites (5 categories). The tree is drawn to scale, with branch lengths measured in the number of substitutions per site. All positions with less than 90% site coverage were eliminated. That is, fewer than 10% alignment gaps, missing data, and ambiguous bases were allowed at any position. Evolutionary analyses were conducted in MEGA7 (Kumar *et al.*, 2016).

### Statistical analyses

The data were analysed using SPSS software (vision 20, IBM). The normal distribution of data and the homogeneity of variances were analysed with the Shapiro-Wilk test (P<0.05) and homogeneity test (P<0.05), respectively. When both tests were not significant, the data were subjected to either one-way or two-way ANOVA followed by a Tukey’s post-hoc test to compare the means of different groups (if there were more than 2 groups). For the data that were either not normally distributed or did not have equal error variances, the data were log or square root transformed and ANOVA analysed on the transformed data.

## Results

### *FAB* and *FIN* are required to suppress arbuscular mycorrhizal development in tomato

Tomato mutants disrupted in *FAB* and *FIN* displayed a significant increase in the mycorrhizal colonisation of the root under the fertilization conditions used, developing approximately 35-50% more total colonisation and arbuscules compared to WT (Fig.1 A, B). For *fab* and both *fin-n2326* lines, this increase in root colonisation was correlated with a significant increase in number of hyphopodia, the fungal entry points along a given length of root, compared to WT (Fig.1 C), suggesting *FAB* and *FIN* may suppress early stages in mycorrhizal development, at or before hyphal entry. Previously published experiments demonstrated that *Slclv2* mutant lines displayed significantly more AM colonisation than WT (Wang *et al.*, 2018). As found previously for *SlCLV2* (Wang *et al.*, 2018), *FAB* and *FIN* appear to influence the amount but not the structure of mycorrhizal features, as the hyphopodia, arbuscules, hyphae and vesicles that formed in mutants disrupted in these genes appeared similar in size and structure to WT (Fig.1 D, data not shown). Phylogenetic analysis using full length amino acid sequences from legume and non-legume families indicate FAB (Solyc04g081590) is related to other CLV1-like proteins, *Sl*CLV2 (Solyc04g056640) is related to *Arabidopsis* and legume CLV2 proteins, and FIN (Solyc11g064850) is related to hydroxyproline *O*-arabinosyltransferase enzymes that influence nodulation in legumes (*Ps*NOD3, *Mt*RDN1 and *Lj*PLENTY; Suppl Fig.S2 at *JXB* online).

**Figure 1.**
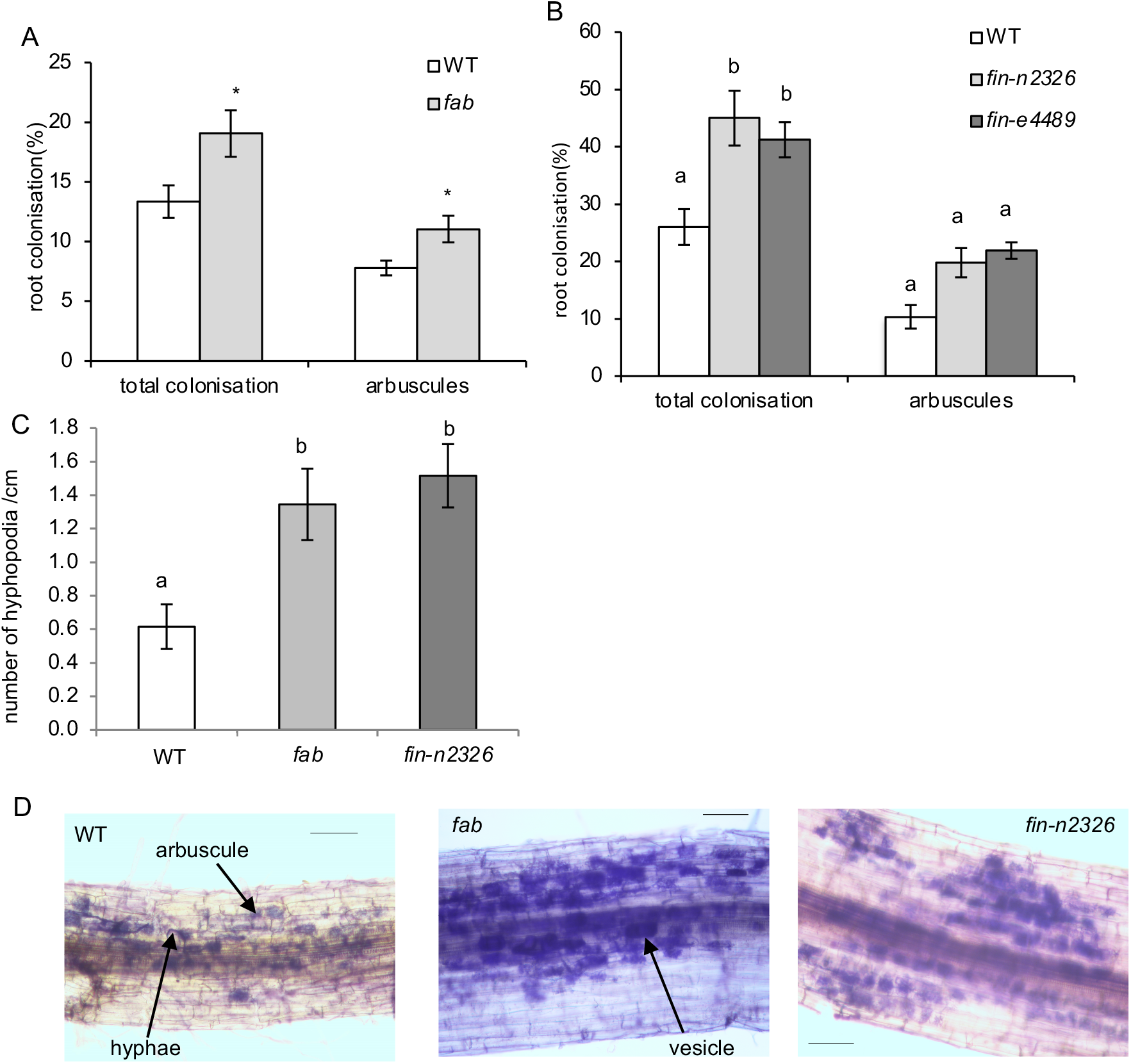
Mycorrhizal phenotype of WT (wild type), *fab* and *fin* mutants grown under 5mM KNO_3_ and 0.5mM NaH_2_PO_4_. (A, B) Percentage of root colonised by any fungal structure and percentage of root colonised with arbuscules, (C) number of hyphopodia per cm of root and (D) photos of typical colonised roots, scale bar is 1mm. (A-C) Data are mean ± standard error (SE) (n=11 - 12 for A, n=6 for B and n= 5 for C). For (A) * indicates values that are significantly different to WT (P<0.05), for (B, C) within a parameter letters indicate values that are significantly different as assessed by Tukey’s HSD test (P < 0.05).

It is important to note that the elevated colonisation of these mutants was not due to increased root or shoot size, as the mutants developed smaller root systems than WT under mycorrhizal conditions (approx. 30-50 % less) and this was mirrored by a similar reduction in shoot size in *fab* and *Slclv2* but not *fin* (Suppl Fig. S3A-D at *JXB* online). Indeed, the intersect scoring method employed in this study should not be influenced by root size or length. A detailed examination of root development of these mutants indicated no significant effects on root architecture of *fab, Slclv2* or *fin* seedlings and only a small reduction in root size in mature non-mycorrhizal *Slclv2* and *fin* mutants compared to WT (Wang *et al.*, 2020b). However, to investigate if root access to inoculum could be responsible for the observed mycorrhizal phenotype, the mycorrhizal colonisation of *fab* mutants and WT plants under two different doses of inoculum was examined (Suppl Fig.S4 at *JXB* online). A two-way ANOVA analysis found a strong genotype effect as expected (P<0.01), but no significant inoculum effect or genotype by inoculum interaction. This suggests that the increased mycorrhizal colonisation rate in *fab* mutants is not simply due to inoculum access under the conditions used.

### Nitrate but not phosphate suppression of mycorrhizal colonisation in tomato requires *FAB* and *FIN*

As has been found for *Petunia hybrida,* rice and *M. truncatula* (Bonneau *et al.*, 2013; Corrêa *et al.*, 2014; Liu *et al.*, 2012; Nouri *et al.*, 2014), application of nitrate significantly suppressed mycorrhizal colonisation in tomato in a dose dependant manner (Fig.2 A,B). In contrast, both shoot and root fresh weight and the shoot:root ratio increased significantly with the increase in nitrate levels (Suppl Fig.S5 A-F at *JXB* online). Clearly, nitrate suppresses mycorrhizal colonisation in tomato, and this suppression is not an indirect effect of low nitrate limiting plant vigour and thus AM formation, as the low nitrate limited plant growth but elevated mycorrhizal colonisation rates.

**Figure 2.**
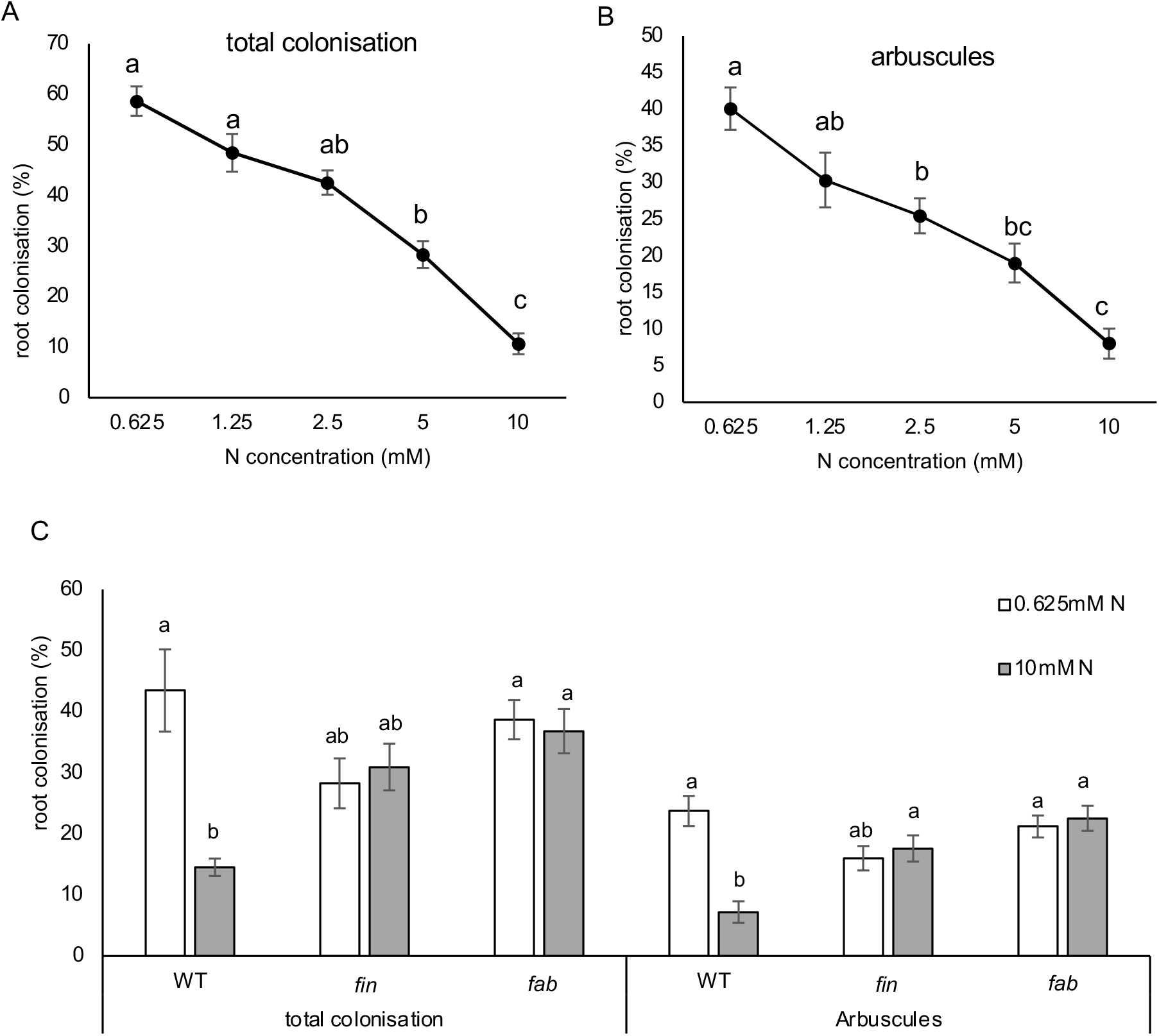
Mycorrhizal colonisation of tomato under various nitrate treatments (0.625-10mM KNO_3_) and 0.5mM NaH_2_PO_4_. (A) Total colonisation and (B) arbuscules of WT (cv. Money Maker) plants grown under different N treatments. (C) WT (cv. M82), *fin-n2326* and *fab* mutants growing under two KNO_3_ treatments (0.625 or 10mM). Data shown are mean ± SE (n=5-6). Within a parameter different letters within a parameter indicate values that are significantly different as assessed by Tukey’s HSD test (P < 0.05).

The mycorrhizal colonisation of WT, *fab* and *fin* plants grown under high and low nitrate conditions was examined (Fig.2 C). As seen in the previous experiment with cv. Money Maker, the total colonisation and arbuscule rate of WT cv. M82 plants growing under high nitrate were both significantly lower than that under low nitrate conditions. Under high nitrate, both *fab* and *fin* mutant plants have significantly elevated arbuscule colonisation compared to WT plants under high nitrate (10mM; Fig. 2C). This is consistent with elevated colonisation seen in *fab* and *fin* plants when grown under relatively high nitrate (5mM; Fig 1). However, an increase in colonisation in response to low nitrate was not observed in *fab* and *fin* mutant plants. In fact, under low nitrate, high rates of colonisation were observed in WT, *fab* and *fin* mutant roots. Two-way ANOVA analysis revealed a strong genotype by treatment interaction (P < 0.001) on both arbuscule and total colonisation, demonstrating that *fab* and *fin* mutants respond differently to nitrate than WT plants. This suggests that *FAB* and *FIN* are required for nitrate suppression of AM and this explains why elevated colonisation is only observed in *fin* and *fab* mutant plants compared to WT when grown under high nitrate (Fig. 1 and 2 C) but not low nitrate (Fig.2 C). As observed in WT, the shoot and root growth of the *fab* and *fin* mutant plants were severely restricted under low nitrate conditions compared to high nitrate (Suppl Fig.S5 D-F). This is consistent with non-mycorrhizal seedling studies, that found that the root development of *fab* and *fin* seedlings responded to altered nitrogen in a similar way to WT (Wanf et al., 2020).

Phosphate is a potent inhibitor of AM colonisation. Indeed, total AM colonisation and arbuscule colonisation of WT roots was reduced more than 10-fold by application of high phosphate compared to WT plants that received low phosphate (Fig.3). This strong suppression of AM colonisation was still observed in *fab* and *fin* mutant plants. Indeed, AM colonisation rates under high phosphate were not significantly different amongst genotypes. Under very low phosphate (0.05mM, Fig.3), a small but not significant increase in arbuscule colonisation rate was seen in *fin* and *fab* mutants compared to WT and this is in contrast to the significant increase in colonisation seen in these mutants when grown under more moderate phosphate limitation (0.5mM; Fig. 1). This suggests that even under relatively high nitrogen (5mM), severe phosphate limitation (0.05mM) strongly promotes mycorrhizal colonisation and this can override the effect of *FIN* or *FAB* on colonisation. Two-way ANOVA analysis revealed a strong treatment effect and small genotype effect but no genotype by treatment interaction for both arbuscule and total colonisation, indicating that *fab* and *fin* mutants respond in a similar way to phosphate as WT plants. As observed in WT, the shoot and root growth of the *fab* and *fin* mutant plants was severely restricted under low phosphate conditions compared to high phosphate (Suppl Fig.S6 at *JXB* online).

**Figure 3.**
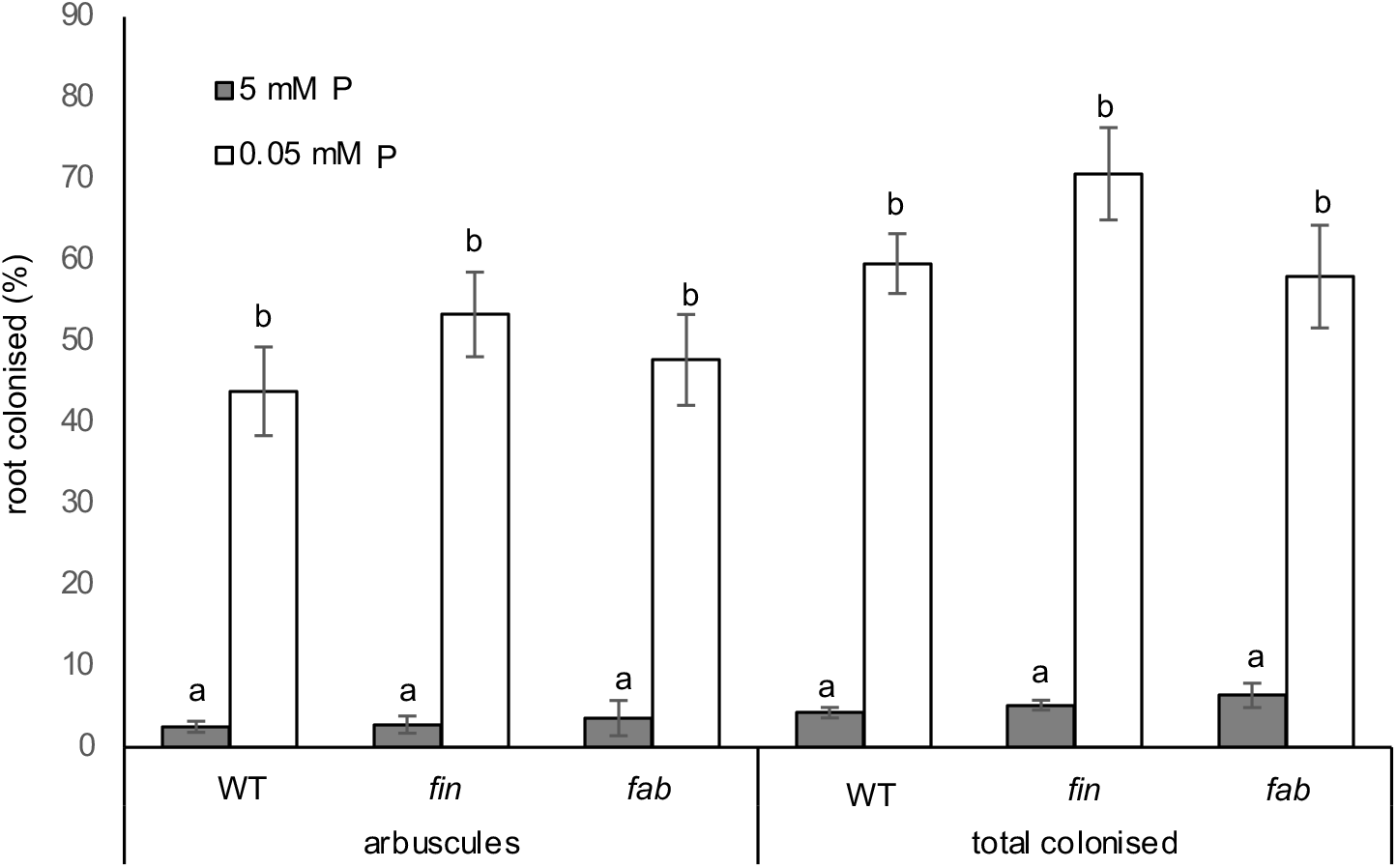
Mycorrhizal colonisation of tomato under low (0.05mM) and high (5mM) phosphate treatments and 5mM KNO_3_. (A) Total colonisation and (B) arbuscules of WT (cv. M82), *fin-n2326* and *fab* mutants growing under two NaH_2_PO_4_ treatments (0.05 or 5mM). Data shown as mean ± standard error (SE) (n=5). Within a parameter different letters indicate values within a parameter that are significantly different as assessed by Tukey’s HSD test (P < 0.05).

### The *FAB* gene acts in the root, while the *SlCLV2* gene may act in both the shoot and root

Grafting and/or split root studies have revealed the *CLV1-like* and *CLV2* genes act in the shoot to suppress nodulation, while *RDN1-like* genes are root acting (e.g. Delves *et al.*, 1986; Sagan, 1996; Schnabel *et al.*, 2011). In addition, there have been at least some studies that suggest *GmNARK* acts in the shoot to suppress AM, although this is not supported by all studies (Meixner *et al.*, 2005; Meixner *et al.*, 2007; Sakamoto and Nohara, 2009). The shoot or root acting nature of *FAB* and *SlCLV2* was tested by reciprocal grafting experiments (Fig.4). As observed in intact plants, colonisation of the root by arbuscules was significantly higher in *fab/fab* (shoot/root stock) self-grafted plants than the WT/WT self-grafts (Fig. 4 A). Importantly, in reciprocal grafts elevated arbuscule rate was only observed when grafts contained *fab* mutant roots. Arbuscule root colonisation was not influenced by shoot genotype, indicating the *FAB* gene appears to act in the root to suppress AM formation. In contrast, only *Slclv2-2/Slclv2-2* self-grafted plants showed significantly higher total colonisation and arbuscule numbers than the WT/WT self-grafted plants (Fig. 4 B). The other graft combinations, WT/*Slclv2-2* and *Slclv2-2*/WT, did not show any significant difference in the extent of colonisation compared with the WT self-graft, suggesting that the *SlCLV2* gene may be acting in both the root and shoot to suppress mycorrhizal colonisation.

**Figure 4.**
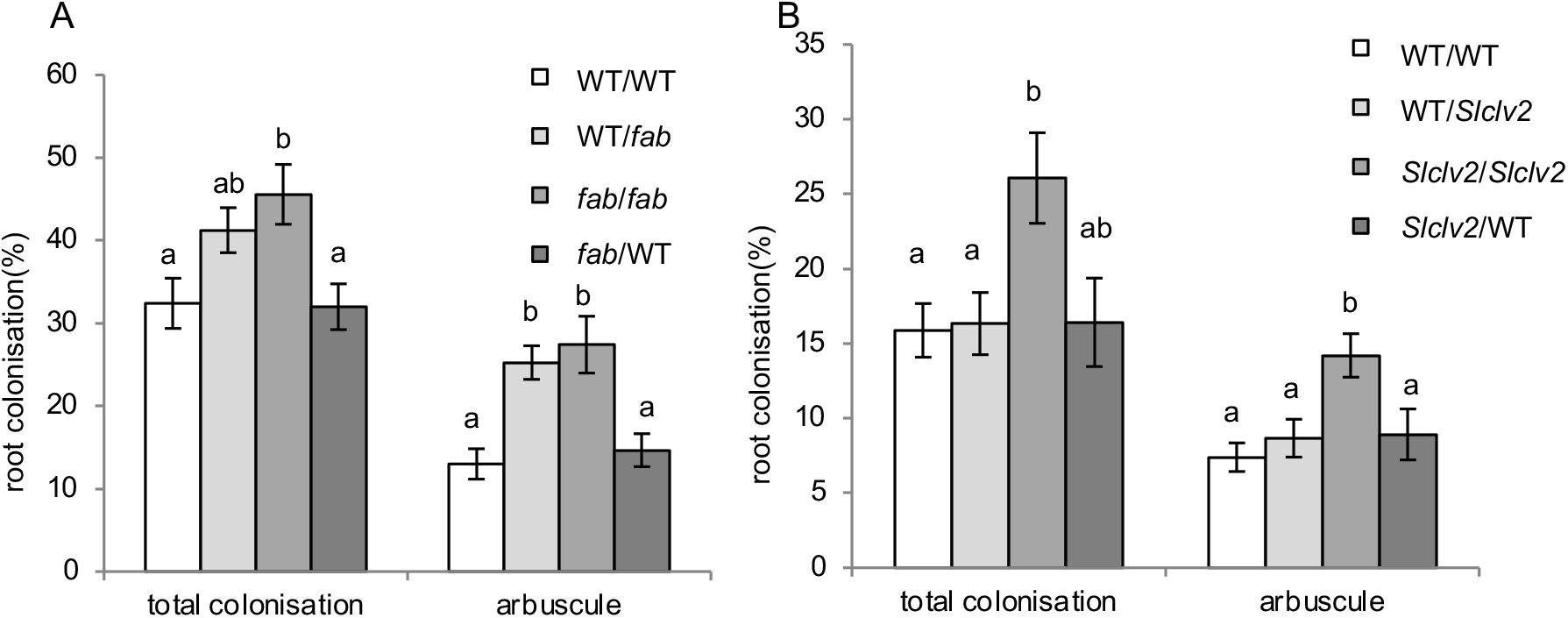
Mycorrhizal colonisation in reciprocal grafts between (A) WT and *fab* and (B) WT and *Slclv2-2* (scion/root stock) grown under 5mM KNO_3_ and 0.5mM NaH_2_PO_4_. Data are shown as mean ± SE (n=8 for *fab* experiment, 8-11 for *Slclv2-2* experiment). Within a parameter different letters indicate values that are significantly different as assessed by Tukey’s HSD test (P < 0.05).

### Strigolactone levels in *fab* and *fin* mutants

Strigolactone levels in the root exudates from the *fab* and *fin* mutants and WT plants were examined under both mycorrhizal colonised and un-colonised conditions (Fig. 5A). Tomatoes produce a variety of strigolactone compounds and orobanchol, orobanchyl acetate and fabacyl acetate were detected in at least some extracts. With one exception, no significant differences in strigolactone levels between mutant and WT plants were observed. The only exception was under non-mycorrhizal conditions, with a small but significant (P<0.05) increase in fabacyl acetate levels in *fin* compared to *fab* and WT. Please note, as the colonised and uncolonised plants were grown at different times they should not be directly compared.

**Figure 5.**
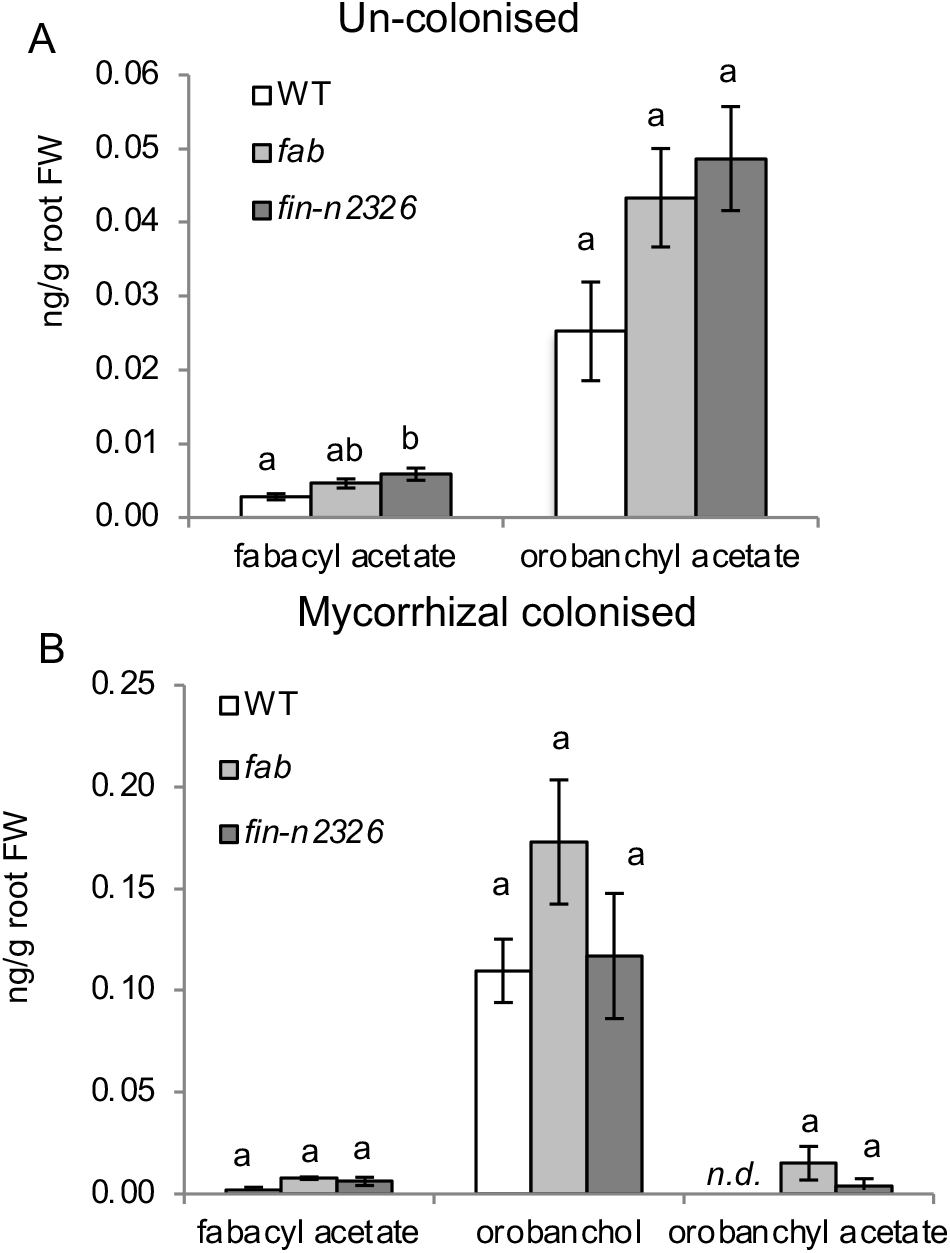
Strigolactone levels in root exudates from tomato WT, *fab* and *fin-n2326* mutants that were colonised by mycorrhizae and grown with 5mM KNO_3_ and 0.5mM NaH_2_PO_4_ (A) or in absence of mycorrhizal inoculum (B). N.B. orobanchol was below detection limit in extracts shown in panel A. Data are shown as mean ± SE (n=5). Different letters indicate values that are significantly different as assessed by Tukey’s HSD test, P < 0.05). *n.d. not detected.*

## Discussion

Studies in this paper suggest roles of an hydroxyproline *O-* arabinosyltransferase enzyme (FIN), an LRR receptor kinase (FAB) and one LRR receptor like protein (*Sl*CLV2) in the negative regulation of AM colonisation in tomato. Nitrate repression of AM also appears to require FAB and FIN, while phosphate appears to influence AM independently of these genes and may override the nitrate response.

The mutant studies indicate suppression of mycorrhizal colonisation of tomato requires the hydroxyproline *O*-arabinosyltransferase enzyme FIN (Fig.1). FIN appears to be important for arabinosylation of CLE peptides active in shoot apical meristem maintenance (Xu *et al.*, 2015). The closest homologs in legumes have been suggested to arabinosylate some CLE peptides essential for AON (Hastwell *et al.*, 2018; Imin *et al.*, 2018; Kassaw *et al.*, 2017; Yoro *et al.*, 2019) and *RDN1* is required for MtCLE53 to suppress AM colonisation in *M. truncatula* (Karlo *et al.*, 2020). Both FAB and *Sl*CLV2 receptors are required to suppress AM symbioses in tomato (Fig.1 and Wang *et al.* (2018)) and grafting indicates FAB appears to act in the root and the shoot and root for *Sl*CLV2 (Fig.4). The role of the *FAB* gene in suppressing mycorrhizal colonisation is consistent with the role of *CLV1-like* genes in the negative regulation of mycorrhizal colonisation of legumes (Meixner *et al.*, 2005; Morandi *et al.*, 2000; Müller *et al.*, 2019; Solaiman *et al.*, 2000). However, it is still unclear if *CLV1-like* genes act in the shoot to suppress mycorrhizae.

Our grafting results suggest *FAB* acts in the root and not the shoot. Similarly in one split root study with soybean *En6500* mutant, disrupted in the *CLV1-like NARK* gene, showed this mutant line retained the ability to systemically suppress AM (Meixner *et al.*, 2007). However, grafting and split root studies with this and other soybean *nark* mutant alleles do suggest *NARK* can act in the shoot to suppress AM (Meixner *et al.*, 2005; Meixner *et al.*, 2007; Sakamoto and Nohara, 2009; Schaarschmidt *et al.*, 2013) indicating a *CLV1-like* gene in soybean may act in the shoot to suppress AM.

As has been found for nitrate regulation of nodulation, in tomato the *CLV1-like FAB* and *RDN1-like FIN* are required for nitrate suppression of AM. Application of high nitrate to tomato suppressed mycorrhizal colonisation and this suppression was absent in *fab* and *fin* mutant plants (Fig.2). This appears to be a specific role for these genes in nitrogen regulation of mycorrhizal colonisation, as these mutants still regulate root architecture in response to nitrogen (NEW ROOT PAPER). This role for FAB and FIN in nitrate response is striking as it suggests deep conservation of the role of these genes in nitrate regulation of symbioses. Future work will explore which other elements of the pathways including which specific CLE(s) might be responsible for this nitrate regulation of AM. In contrast, the very strong suppression of AM colonisation by high phosphate did not require FAB and FIN, as the suppression of AM by phosphate was maintained in lines disrupted in these genes (Fig.3). This is also consistent with studies in homologous legume mutants, which found that phosphate suppression of AM was not disrupted in *clv1-like* mutants in pea, *M. truncatula* and soybean (Foo, 2014; Müller *et al.*, 2019; Wyss *et al.*, 1990). Thus, phosphate regulates AM via pathway(s) independent of CLV1-like proteins across species and FIN in tomato. Given the suite of CLE peptides, modifying enzymes and receptors in each plant, the possibility that particular stimuli (e.g. symbioses, nitrate, phosphate, etc) acting through specific combinations of signals and receptors is likely.

Strigolactones are at least one, but not the only signal through which phosphate suppresses AM (Balzergue *et al.*, 2011; Breuillin *et al.*, 2010; Foo *et al.*, 2013b). It is difficult to distinguish if strigolactones are acting not only as an upstream signal but also as a downstream signal in the AOM pathway. This is because strigolactones are essential for substantial colonisation to occur and it is clear that a certain level of colonisation is required to induce AOM (Vierheilig, 2004). It is intriguing that the *FAB* and *RDN1* genes appear to suppress colonisation in part through suppressing fungal entry and this is consistent with studies in *M. truncatula* with *CLE53/33* overexpression lines (Müller *et al.*, 2019). However, strigolactones levels are not elevated in *clv1-like* mutants of pea, *M. truncutula* or tomato or *RDN1-like* mutants in pea and tomato (Foo *et al.*, 2014; Müller *et al.*, 2019), meaning elevated strigolactone exudation does not appear to be the cause of elevated AM colonisation of these mutants. Clearly, further clarification of the interaction between strigolactones and these genes is required.

Grafting studies presented here suggest in tomato *SlCLV2* can act in the shoot and root to suppress AM. This is in contrast to the clear role for this gene in the shoot but not the root during AON of legumes. However, as the single study in pea examining the role of *CLV2* in AM regulation of legumes found no influence of this gene on AM formation (Morandi *et al.*, 2000), further studies are required to clarify the action of *CLV2* in AM regulation of legumes and non-legumes. It is important to note that split root studies indicated that negative regulation of AM can act systemically (Vierheilig *et al.*, 2000). Therefore, it will be important to clarify which genes and signals may mediate this long-distance effect and which may act locally in the root to limit AM formation. Studies with *M. truncutula* plants indicated that although *MtCLE53* expression was induced by AM locally, a systemic effect of this peptide on AM was suggested by transgenic studies with chimeric roots overexpressing *MtCLE53* (Karlo *et al.*, 2020). However, further studies are required to clarify the tissue(s) in which other elements of the pathway act. Future studies could also begin to explore the nature of the receptor complex(es) that may be important for negative regulation of AM by specific CLE peptides, including examining if CLV1-like and CLV2 proteins work in parallel or as co-receptors and the role of other receptors that play roles in AON (e.g. KLV and CRN). Indeed, recent research has shown that compensation mechanisms operate in the CLE - CLV signalling pathway to control shoot apical meristem homeostasis through both ligands and receptors (Nimchuk *et al.*, 2015; Rodriguez-Leal *et al.*, 2019).

Work presented in this paper enables us to build a model of the genetic regulation of mycorrhizal colonization in a non-legume tomato. In addition to roles for *CLV1-* and *RDN1*-like genes previously identified in legumes (and for a *CLV1-like* gene in *Brachypodium),* this model indicates these genes are important for nitrate but not phosphate regulation of mycorrhizal colonization and includes an important role for *SlCLV2* (Fig. 6). Specific CLE peptides have been identified to act via SUNN and RDN1 in *M.truncutula* (Karlo *et al.*, 2020; Müller *et al.*, 2019), and future studies will examine whether similar CLE peptides influence AM in tomato. The expansion of our understanding of the negative regulation of AM into non-legume systems and the parallels this pathway shares with AON in legumes, including nitrate regulation, suggests that negative regulation of symbioses is a conserved symbiotic program. In addition to the common symbiotic pathway essential for early signaling and infection events in nodulation and AM (Radhakrishnan *et al.*, 2020), we might consider this negative regulation of symbioses via autoregulation as the bookend to this pathway and therefore a component of the so called ‘common symbiotic pathway’(Chiu and Paszkowski, 2020). Indeed, recent studies have suggested AON may influence a subset of genes, including the Nod-factor receptor NFP (Gautrat *et al.*, 2019), integrating these two programs. It is also interesting to consider that although the CLV-CLE pathway shares an orthologous role in shoot apical meristem control in non-legumes, no clear role for *CLV1-*like genes in shoot development of legumes has been observed (e.g. Krusell *et al.*, 2002; Nishimura *et al.*, 2002). Thus, an orthologous role for CLV1-like proteins in suppression of AM in both legumes and non-legumes (tomato this paper, and *Brachypodium* (Müller *et al.*, 2019)) describes a common function for this gene across legumes and non-legumes. A deeper understanding of how plants manage interactions with nutrient acquiring microbes provides a basis for future advances in plants acquiring maximum benefit from these symbiotic relationships.

**Figure 6.**
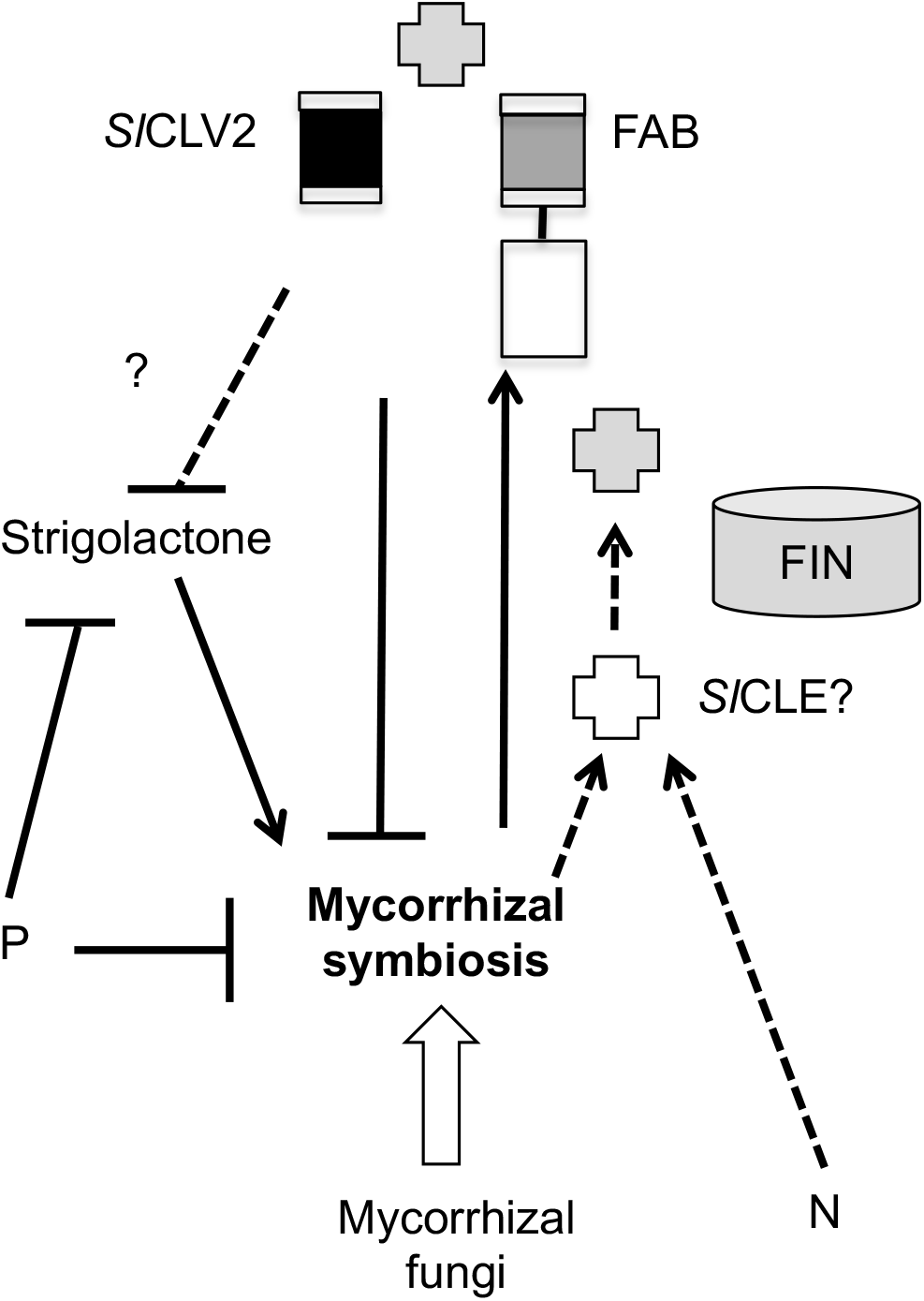
A model of the action of FAB, *Sl*CLV2 and FIN on mycorrhizal colonization of tomato, including the influence of nitrate (N) and phosphate (P). Flat-ended lines indicate a negative influence, while arrows indicate a positive influence. Question marks and dotted lines indicate unclear or as yet untested elements.

## Supplementary data

**Suppl Figure 1.** Outline of CRISPR guide RNA and mutations introduced in *Slclv2-2* lines.

**Suppl Figure 2.** Phylogenetic tree of (A) CLV1-like, (B) CLV2, and (C) HPAT family proteins.

**Suppl Figure 3**. Growth parameters of WT (cv. M82), *fab, Slclv2 and fin* mutants grown with mycorrhizal inoculum as shown in Fig. 1 for *fab* and *fin* and from Wang *et al.* (2018) for *Slclv2.*

**Suppl Figure 4**. Mycorrhizal colonisation of WT and *fab* plants mutants grown with standard (20%) or high (40%) inoculum doses.

**Suppl Figure 5.** Growth parameters of WT (cv. Money Maker), WT (cv. M82), *fin-n2326* and *fab* mutants grown under various KNO_3_ treatments.

**Suppl Figure 6.** Growth parameters of WT (cv. M82), *fin-n2326* and *fab* mutants growing under two NaH_2_PO_4_ treatments (0.05mM or 5mM).

## Data availability statement

All data supporting the findings of this study are available within the paper and within its supplementary materials published online.

## Acknowledgements

We sincerely thank Prof Zachary Lippman (Cold Spring Harbour, USA) for his kind gift of tomato mutant lines and for helpful discussions. Many thanks to Prof Koichi Yoneyama (Utsunomiya University) and Dr Kohki Akiyama (Osaka Prefecture University) for their kind gifts of measured quantities of labeled strigolactones and to A/Prof Chris McErlean (University of Sydney) and Dr Bart Janssen (The New Zealand Institute for Plant & Food Research Limited) for the gift of unlabeled solonocal. We thank Michelle Lang, Tracey Winterbottom, Shelley Urquhart and Valerie Hecht (University of Tasmania) for technical advice and assistance.

E.F. was supported in part through the ARC Future Fellowship FT140100770, the work carried out in Australia was funded through the ARC Discovery project DP140101709 and C.W. and K.W. were supported by a Tasmania Graduate Research Scholarship and an Australian Research Training Program scholarship respectively. C-T.K. was supported by the National Science Foundation Plant Genome Research Program (grant no. IOS-1546837).

## List of author contributions

E.F. conceived the project; C-T.K. generated the CRISPR lines; E.F. and J.B.R. supervised the other experiments; C.W. performed the experiments and analyses except the phosphate experiment that was performed by K.W; D.S.N. developed the UPLC/M-MS method; K.V.P. provided technical assistance; E.F., C.W. and J.B.R. wrote the manuscript. E.F. agrees to serve as the author responsible for contact and ensures communication.

