## Supplementary material for "The role of CLV signalling in the negative regulation of mycorrhizal colonisation and nitrogen response of tomato": Suppl Figures

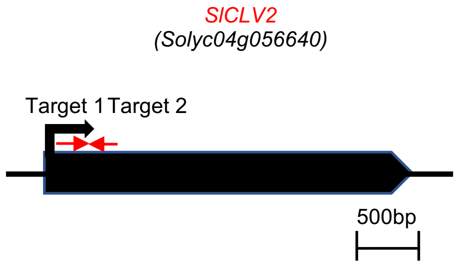


*SlCLV2* GGGTGTTCAATCTTGAAACAGAAG**<u>AGG</u>**ACATGGG (55) GTTAGTGTTCCTCTGTCAAACTGGAC**<u>TGG</u>**TG
*Slclv2-2* GGGTGTT----------------G**<u>AGG</u>**ACATGGG (55) GTTAGTGTTCCTCTGTCAAACTGG----**<u>G</u>**TG

**Suppl Figure 1.** Outline of CRISPR guide RNA and mutations introduced in *Slclv2-2* (change in amino acid sequence after valine (74) and premature stop codon (83)).

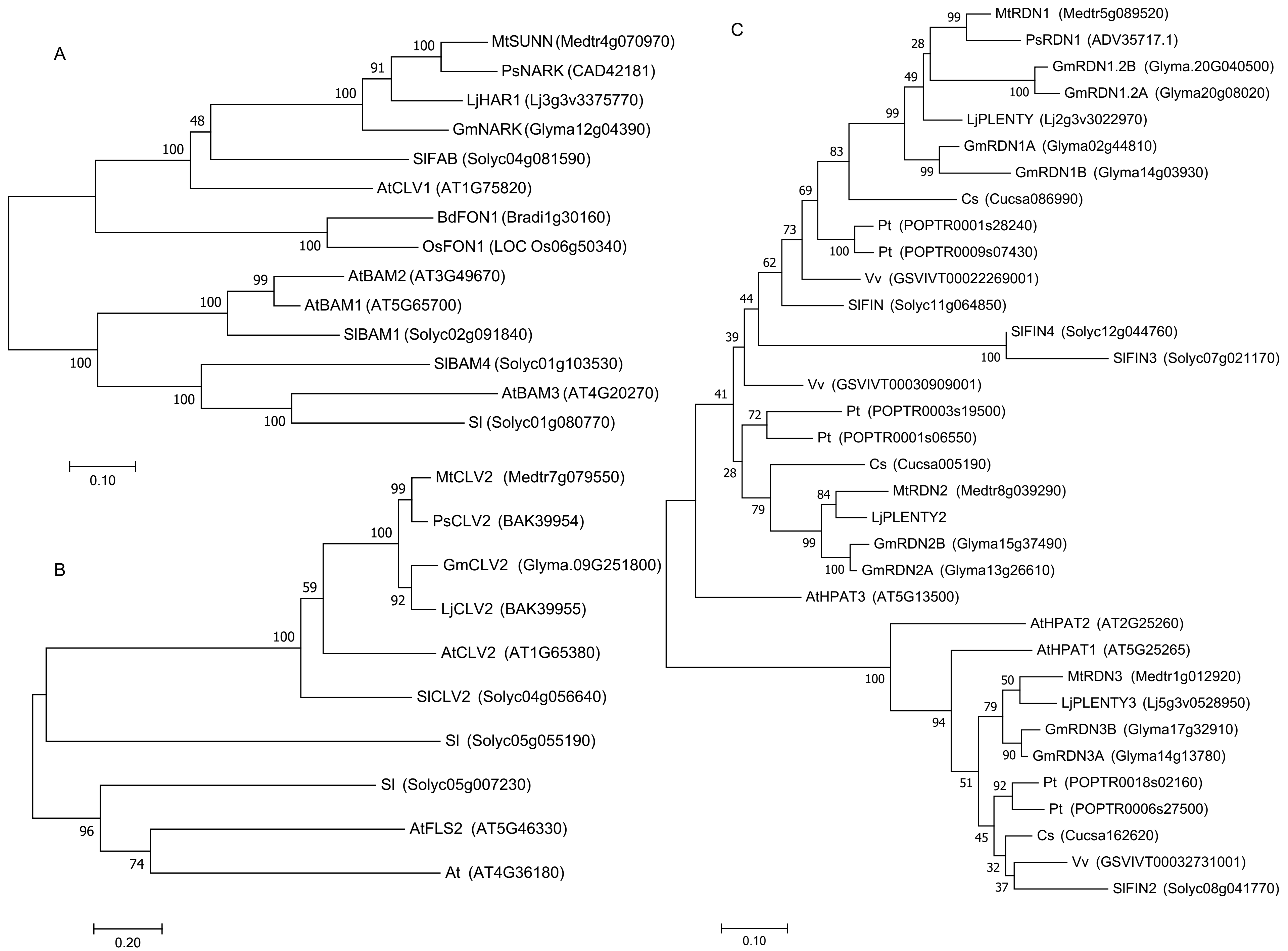


**Suppl Figure 2. Phylogenetic tree of (A) CLV1-like, (B) CLV2, and (C) HPAT family proteins.**

The sequences included are from tomato *(Solanum lycopersicum, Sl), Lotus japonicus* (Lj), *Medicago truncatula* (Mt), soybean (*Glycine max*, Gm), grape (*Vitus vinifera*, Vv), cucumber (*Cucumus sativus*, Cs), poplar (*Populus trichocarpa*, Pt), *Brachpodium distachyon* (*Bd*), *Oryza sativa* (Os) and *Arabidopsis thaliana* (At). Bootstrap values are indicated.

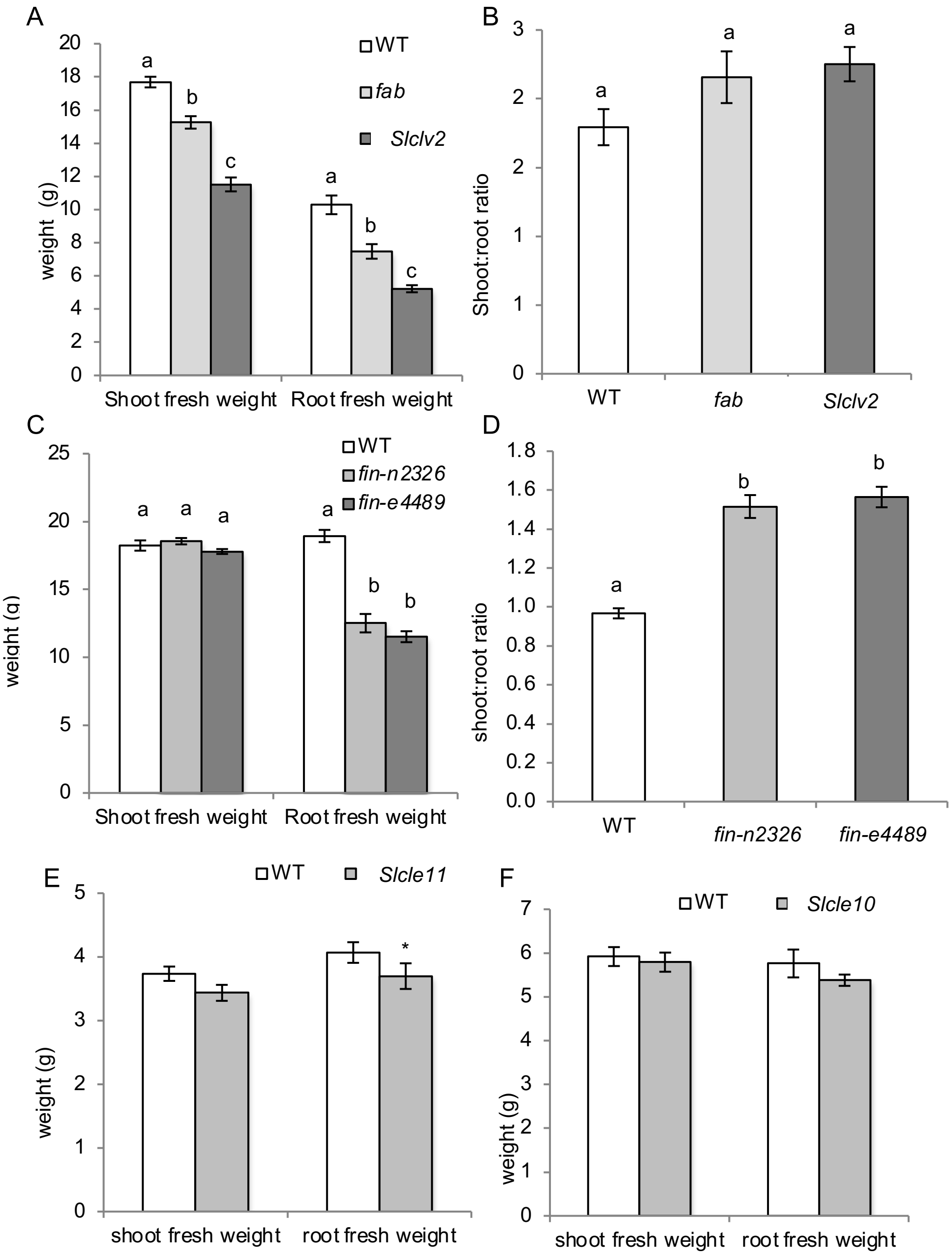


**Suppl Figure 3**. Growth parameters of WT (cv. M82), *fab, Slclv2 and fin* mutants grown with mycorrhizal inoculum and 5mM $KNO_3$ and 0.5mM $NaH_2PO_4$ as shown in Fig 1 for *fab* and *fin* and from Wang et al., (2018) for *Slclv2*. Data are shown are mean ± standard error (SE) (n=12 -13). Within a parameter different letters indicate values that are significantly different as assessed by Tukey's HSD test (P < 0.05).

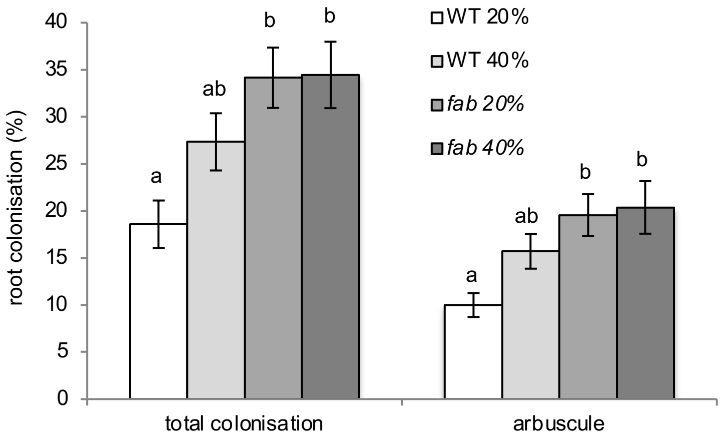


**Suppl Figure 4**. Mycorrhizal colonisation of WT and *fab* plants mutants grown with standard (20%) or high (40%) inoculum doses and 5 mM $KNO_3$ and 0.5 mM $NaH_2PO_4$. Data are shown as mean ± standard error (SE) (n=8 - 9). Within a parameter different letters indicate values that are significantly different as assessed by Tukey's HSD test ($P < 0.05$).

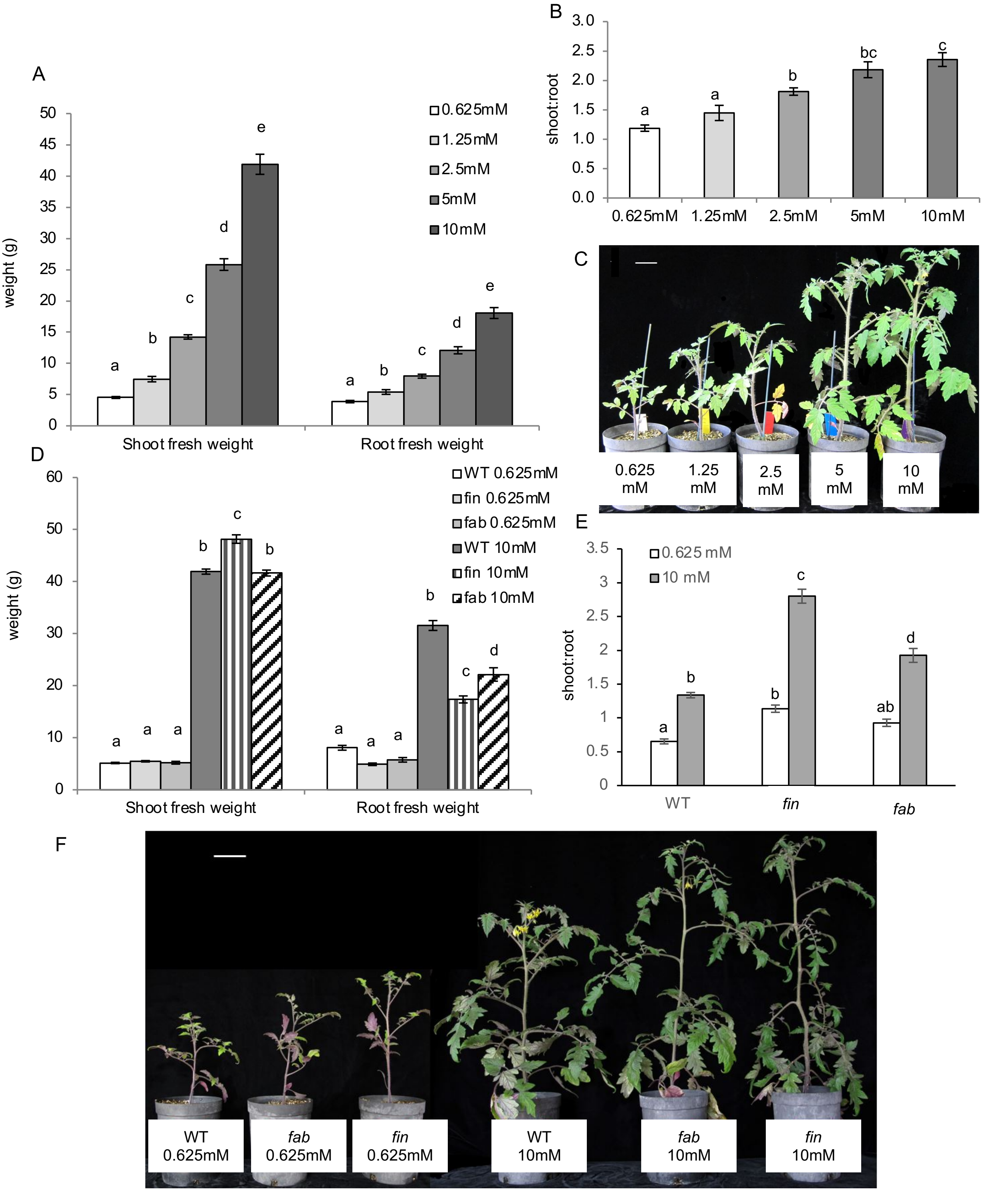


**Suppl Figure 5.** Growth parameters of WT (cv. Money Maker), WT (cv. M82), *fin-n2326* and *fab* mutants grown under various nitrate treatments (0.625-10mM $KNO_3$) and 0.5mM $NaH_2PO_4$. (A-D) Shoot and root fresh weight, shoot:root ratio and photo of WT (cv. MoneyMaker) and (E-G) shoot and root fresh weight, shoot:root ratio and photo of WT (cv. M82), *fin-n2326* and *fab* mutants. Data are shown as mean ± SE, (n=10-12). Values with different letters within a parameter indicate values that are significantly different as assessed by Tukey's HSD test ($P < 0.05$). Scale bar = 5cm.

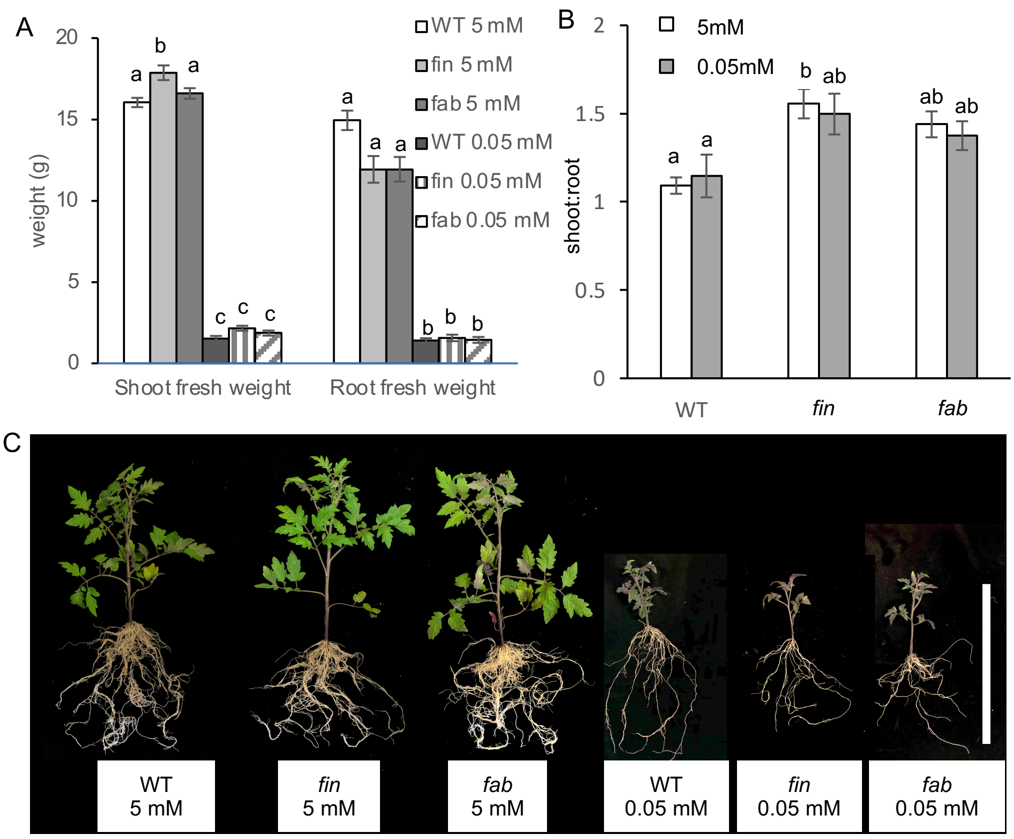


**Suppl Figure 6.** Growth parameters of WT (cv. M82), *fin-n2326* and *fab* mutants grown under low (0.05mM) or high (5mM) phosphate treatments and 5mM $KNO_3$ (A) Shoot and root fresh weight, (B) shoot:root ratio and (C) photo of shoot phenotype, scale bar is 30cm. Data shown are mean ± standard error (SE) (n=11-12). Different letters indicate values within a parameter that are significantly different as assessed by Tukey's HSD test ($P < 0.05$).
